# A Universal Algorithm to Detect Rare or Novel Cell Types in High-Throughput Single-Cell Gene Expression Data

**DOI:** 10.1101/438382

**Authors:** Zoe Weiss, Daphne Tsoucas

## Abstract

Detecting rare cell types would allow early disease detection of cancers and infections, identification of new cell types, and a deepened understanding of cell differentiation. We developed a universal algorithm to identify rare cell types from a wide variety of single-cell ’omics’ data. We validated our algorithm on single-cell qPCR data from mouse hematopoietic cells and single-cell RNA-seq data from human glioblastoma tumors cells, both with expression values from an ample number of genes. We then applied our algorithm to seq-FISH data from mouse hippocampus cells containing expression values from only 121 genes. Our algorithm detected rare cell types including a putative new hippocampal cell type.

**Author summary:** Rare cell type detection would advance early disease diagnosis (e.g., cancer, infection), allow identification of new cell types, and increase understanding of cell differentiation. Current computational methods can detect common cell types, but it remains a challenge to detect rare cell types within cell populations, especially with expression data from a relatively small number of genes. We created a powerful algorithm to detect rare cell types in a population of cells. We validated our algorithm on data from mouse blood stem cells and human glioblastoma tumors cells. When we applied our algorithm to mouse hippocampus data containing expression values from only 121 genes, we detected a putative new brain cell type that no previous algorithm has identified. Our universal algorithm can now be applied to a wide range of data to detect the early onset of diseases and discover new cell types.

## Introduction

Studying single-cell ’omics’ provides fine scale analysis and discovery of distinctive features not seen in cell population averages [1]. A current challenge in single-cell analysis is detecting rare cell types [2, 3]. Known cell types that play a key role in development or disease often have low frequencies in the human body [4]. Some rare cell types include circulating tumor cells (CTCs) and cancer stem cells [5]. The frequency of CTCs is 1 in 10^9^ blood cells, and cancer stem cells are < 1% of total tumor cell population [6]. Because they are so rare, they are often undetected or considered outliers [7]. Detecting rare cell types would allow early disease detection (e.g., cancer, infection), identification of new cell types, and new insights into cell differentiation [2, 8].

GiniClust, the state-of-the-art computational method to detect rare cell types, uses a ”Gini index scale” that compares the expression level of analogous genes in a population [11]. If a cluster of cells expresses these genes differently than most other cells, but similarly to each other, it is classified as a rare cell type. GiniClust has demonstrated a high sensitivity and specificity to detect rare cell types in large cell populations. However, GiniClust is unable to analyze data with gene expression values from a small number of genes, e.g. seq-FISH. In addition, GiniClust can not detect large clusters containing different cell types, which may contain rare (or new) cell types. GiniClust selected genes are not necessarily differentially expressed between major cell clusters. This particular issue has been partly addressed by GiniClust 2 [9].

We developed a new, universal algorithm which overcomes these limitations and can be applied to a wider range of ’omics’ data [10]. Applying our method to mouse hippocampus seq-FISH data, we discovered a putative new type of hippocampal cell.

## Methods Overview

Existing methods to detect rare cell types from ’omics’ data work well on qPCR and RNA-seq datasets: data types with expression values from an ample number of genes. With the goal of analyzing other types of ’omics’ data (e.g. seq-FISH data with expression values from relatively few genes), we developed a two-step cell classification method to identify rare cells based on two novel indices: cell homogeneity and gene homogeneity. The first step is to use gene expression values to classify cells into large clusters and the second step is to detect rare cell types within each large cluster. We view the first step as filtering for the second step.

More specifically, the cells are first clustered into large cell groups based on the similarity of their gene expression using all genes. Within these large groups, we distinguish cells with high cell homogeneity indices. Using gene expression values from these cells, we select the genes that have the highest gene homogeneity indices. We identify rare cell types using the machine learning clustering algorithm Density-Based Spatial Clustering of Applications with Noise (DBSCAN), which utilizes the gene expression values for high homogeneity index genes (Figure 1).

**Fig 1.**
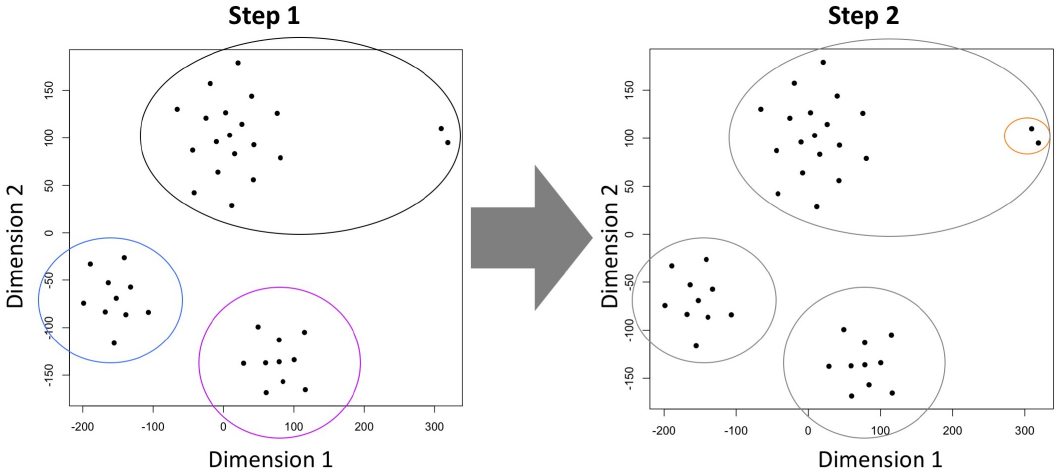
Single-cell gene expression data are plotted using tSNE, a machine learning tool to visualize high-dimensional data. The subfigures depict the two steps of this method. The first step finds large clusters (black, purple, and blue on the left), and the second step finds rare cell types (orange on the right). Our algorithm detects one rare cell type, which is circled in orange.

## Results

### 0.1 Mouse Hematopoietic Cell qPCR Dataset

We first tested our new algorithm on a single cell data from 1963 mouse hematopoietic cells with expression values from 300 genes (Figure 2) [11]. Our method identified two rare cell clusters comprised of 24 mammary gland stem cells and 23 intestinal stem cells, based on their expression values for 74 genes. We visualized the clusters with tSNE, which indicates that the rare cell types are distinct from each other.

**Fig 2.**
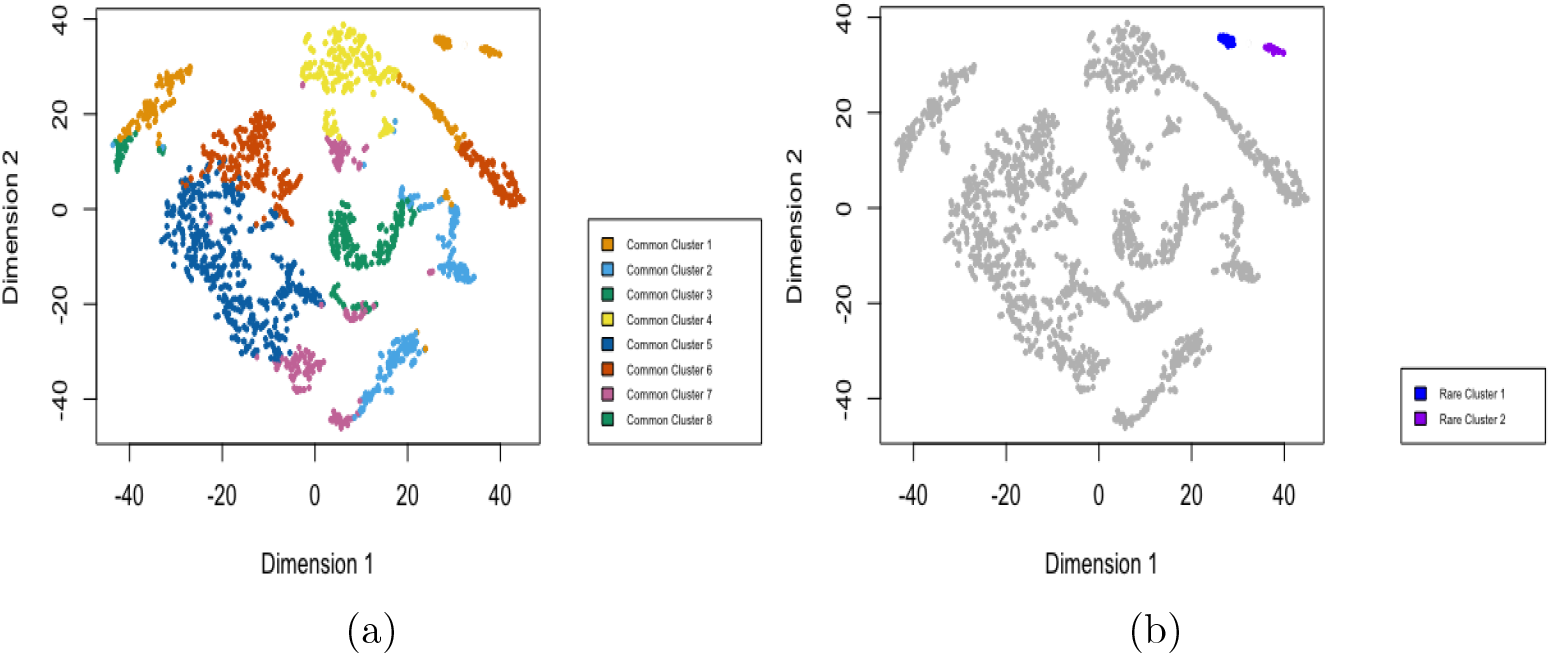
(a) Step 1: Large Group Clustering. tSNE plot of output from the first step of our algorithm applied to single-cell mouse hematopoietic cell data from 1963 cells. (b) Step 2: Identify Rare Cells Within Clusters. This dataset contains two rare cell types, mammary gland stem cells (MASC) and 23 intestinal stem cells (ISC). Our method clearly identified the two rare cell types and differentiated between them. The rare cell types are highlighted in dark blue and purple [11].

We then compared these results to those of GiniClust where the identified rare cell types matched completely. This provided the first validation of the discrimination properties of our method [12]. In the next two sections, we compared our method with GiniClust in other datasets, where the cell groups were more heterogeneous.

### 0.2 Human Glioblastoma Primary Tumor Cells RNA-seq Dataset

We next analyzed a single-cell RNA-seq dataset consisting of 576 human glioblastoma (GBM) primary tumor cells with expression values from 23,234 genes. Gene expression of tumor cells is significantly more heterogeneous than blood cells, so this provides a tougher test of our method [13, 14].

Our algorithm identified 34 high homogeneity index genes and 29 rare cells (Figure 3). GiniClust identified 10 of these cells. The discrepancy is likely due to the different thresholds for the number of genes utilized between GiniClust and our method.

**Fig 3.**
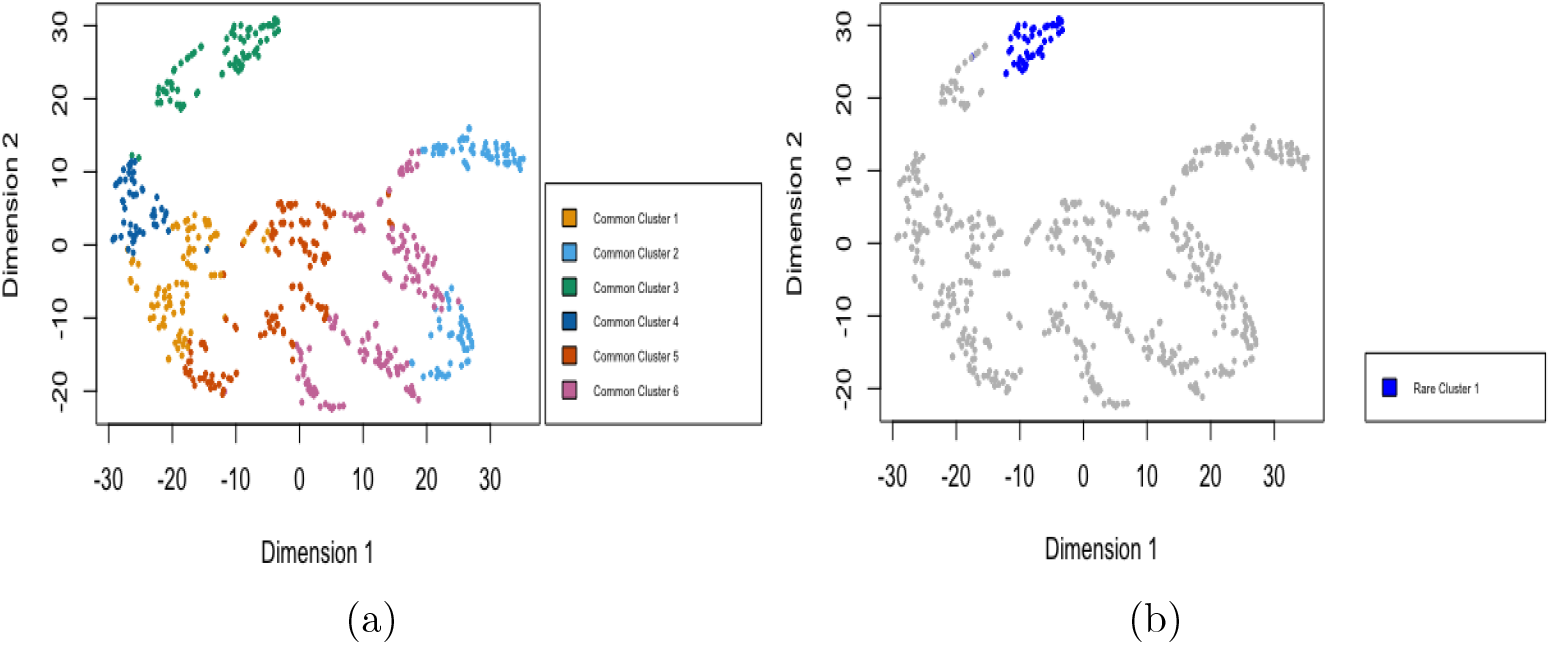
(a) Step 1: Large Group Clustering. tSNE plot of output from the first step of algorithm applied to single-cell human glioblastoma primary tumor cell data from 576 cells. (b) Step 2: Identify Rare Cells Within Clusters. This dataset contained one rare cell type, which our method accurately identified. The rare cell type is highlighted in dark blue [11].

### 0.3 Mouse Hippocampus seq-FISH Dataset and a putative new cell type

Finally, we analyzed a seq-FISH dataset consisting of 2586 cells obtained from a mouse hippocampus with expression values from 121 genes (Figure 4) [15]. Notice that there are far fewer genes than the previous examples. Existing algorithms do not identify any rare cell types. Our algorithm selected 29 genes with high variation between cells, which included four cell types unrecognized by GiniClust. Our method can detect rare cell types with a small number of genes.

**Fig 4.**
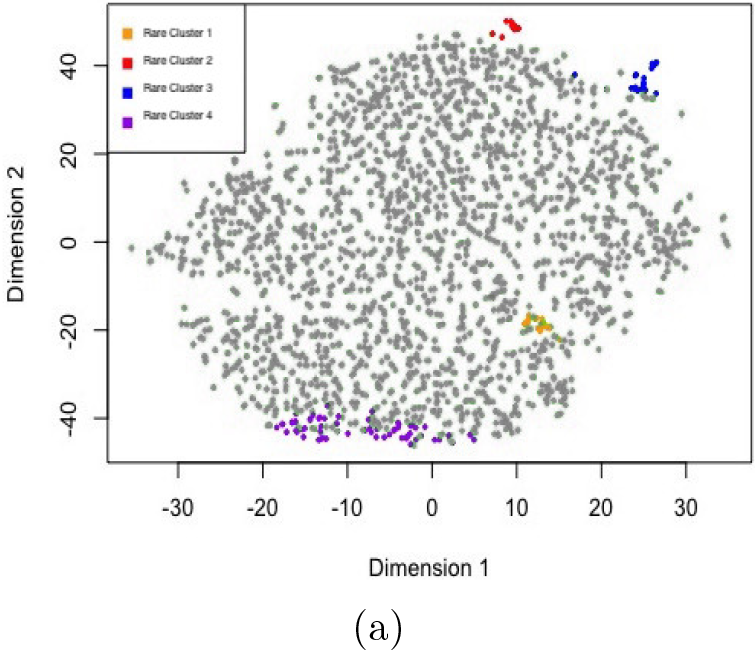
tSNE plot of output from our algorithm applied to single-cell hippocampus data from 2586 cells. Our method identified four previously unidentified groups of transcriptionally distinct cells, which may be biologically significant. The rare cell types are highlighted in orange, purple, blue, and red, with outliers highlighted in pink [15].

We employed the Moran’s I spatial correlation statistic to quantify the spatial correlation of the four detected rare cell types, see Materials and Methods section. This index is normalized to lie between −1 and 1. A high Moran’s I correlation coefficient indicates that the cells are highly correlated in space and in gene expression.

The Moran’s I correlation coefficients for Cluster 1 and Cluster 2 were 0.83 and 0.82, with statistically significant p-values (Table 1). This indicates that the cells in the novel cell groups are highly clustered. Cluster 3 and Cluster 4 had low Moran’s I correlation coefficients. This indicates that the cells in these clusters are likely outliers and not a rare cell type.

**Table 1.** Moran’s I correlation coefficients

| Cluster | Moran's I | p-value | E(Moran's I: H0) | Standard Deviation |
| --- | --- | --- | --- | --- |
| 1 | 0.83046 | 6.32E-13 | -0.00472 | 0.0341 |
| 2 | 0.82289 | 3.00E-14 | -0.025 | 0.11386 |
| 3 | 0.25397 | 0.03296 | -0.14285 | 0.18609 |
| 4 | 0.13889 | 0.16385 | -0.33333 | 0.33919 |

To investigate whether the cells in Cluster 1 and Cluster 2 are actual cell types, we searched four cell atlases for known cell types with similar gene expression: Harmonizome, Allen Brain Atlas, mousebrain.org, and European Bioinformatics Institute [16–19].

Cluster 2 registered as a mast cell type; a mast cell expressed the seven of the ten highest expressed genes from Cluster 2. No known cell types expressed more than three of the ten highest expressed genes from Cluster 1. Thus, Cluster 1 appears to be a putative new cell type.

The top genes expressed in Cluster 1 are *acta2, cldn5, col5a1, dcx, gda, loxl1, nes, rhob, slc5a7,* and *sox2*. We investigated the known functions of these genes individually. We hypothesize that this putative cell may be a new type of neural stem cell with enrichment for genes associated with growth and movement.

## Conclusion

Single-cell analytics has transformed the way we study cell populations and could allow for early disease detection, extend our knowledge about transcriptional mechanisms behind tissue differentiation processes, and allow us to identify new cell types [20]. A major challenge is to detect rare cell types. We developed a universal machine learning based algorithm to identify cell types using single-cell ’omics’ data. We validated it on qPCR and RNA-seq datasets (with an ample number of genes expressed) and obtained similar results to GiniClust and GiniClust 2 [21]. We then applied the algorithm to a seq-FISH dataset containing only 121 genes and our method discovered a putative new type of hippocampal cell.

To compare our method with the state-of-the-art algorithm, GiniClust quantifies the inequality of gene expression with a ”Gini Index scale” in a single step and selects the cells containing genes expressed with the highest ”Gini index”. Our new method first compares expression distances between cells and clusters them into large clusters. Next it identifies potential rare cells within these clusters. We view this first step as filtering. It then clusters the potential rare cells into rare cell types based on their expression of high homogeneity index genes. GiniClust was not designed to detect large clusters which may contain rare (or new) cell types, because the genes that GiniClust selects need not be differentially expressed genes between major cell clusters, as in the third dataset.

Our new, universal algorithm which overcomes these limitations can be applied to a wider range of ’omics’ data.

## Materials and methods

### 0.4 Data

We validated our algorithm on two datasets: 1) qPCR from 280 mouse hematopoietic cells with expression values from 300 genes [11], 2) RNA-seq from 576 human glioblastoma (GBM) primary tumors cells with expression values from 23,234 genes [11]. We then applied our algorithm to seq-FISH from 2586 mouse hippocampus cells with expression values from only 121 genes [15].

### 0.5 Step 1: Large Group Clustering

#### 0.5.1 Preparation of Data

In some single-cell data, e.g RNA-seq, zero values are common. To increase the robustness of our algorithm, we first remove genes with zero or negligible expression and then normalizes the remaining data to have unit variance. The excised data were either data collection errors or not regularly expressed genes.

#### 0.5.2 Defining Cell Distances

We define a distance between cells using the non-parametric Spearman correlation. Correlations measure similarity between cells. The closer the correlation is to one, the greater the association is between the cells. However, this method is based on quantifying dissimilarity between cells, so we convert correlation to distance. Distance is defined as: Distance = 1 − Spearman Correlation

This transformation is applied to the entire Spearman correlation matrix, outputting a new matrix with dissimilarity between each pair of cells.

#### 0.5.3 Similar Cell Clusters

We use k-means clustering to cluster cells into smaller, more similar groups based on the distance between them. k-means requires the number of clusters, k, and the distance matrix as inputs. In general, there is no robust algorithm to optimize k. However, we discovered the following unexpected linear relationship:

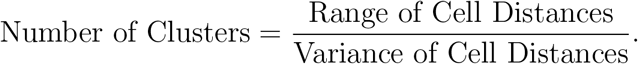

### 0.6 Step 2: Identify Rare Cells Within Clusters

We assign each cell within a large cluster a novel cell homogeneity index (Cell HI) summing the variance from the mean gene expression for each gene expression value.

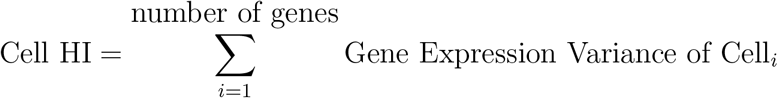

where Gene Expression Variance of Cell_*i*_ is the variance of the gene expression values from each gene for a given cell compared to the the mean gene expression values for that cell.

The cells with homogeneity indices far from the mean are considered potential rare cells. For each potential rare cell, we select the genes with the highest gene homogeneity indices (Gene HI), identified as a clear threshold above the mean.

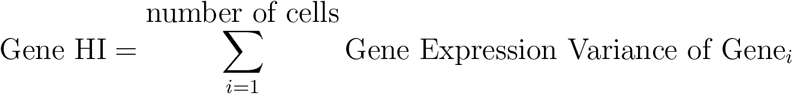

where Gene Expression Variance of Gene_*i*_ is the variance of the gene expression values from each cell for a given gene compared to the the mean gene expression value for that gene. We trim the original data to only retain potential rare cells and genes with high homogeneity indices.

#### 0.6.1 DBSCAN

To identify the putative rare cell types within the potential rare cells (all rare cell types and outliers), we use the clustering algorithm, DBSCAN (from the ’dbscan’ package in R). We use DBSCAN, instead of *k*-means, because it does not require the number of clusters as an input [22]. DBSCAN requires two parameters: the minimum number of cells to be considered a cluster, and ε being the minimum distance between two cells to be grouped in the same cluster. There is no computational method to determine ε, so we test multiple values. This method outputs how many rare cell types are present, and which cells belong to which rare cell type.

#### 0.6.2 Visualize Using tSNE And Compare To GiniClust

We use t-distributed stochastic neighbor embedding (tSNE), to visualize our very high-dimensional data. We compare the findings of our method to those of the current standard, GiniClust.

#### 0.6.3 Compute Moran’s I Spatial Correlation

Moran’s I measures spatial correlation based on both feature locations and values simultaneously. Up until now, we could not determine if our putative rare cells were spatially clustered. For this, we employ Moran’s I spatial correlation:

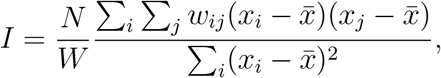

where *N* is the total number of rare cells (indexed by *i* and *j*), *x* is the gene expression, *x*̄ is the mean of *x*, *w_ij_* is the matrix of the squares of the physical distances between cells, *W* is the sum of all *w_ij_*.

A Moran’s I value close to 1 for a rare cell group signifies a very strong spatial correlation.

#### 0.6.4 Compare to Known Cell Types

Our method identifies a few potential new cell types. We search cell atlases (Harmonizome, Allen Brain Atlas, mousebrain.org, and European Bioinformatics Institute for all known brain cell types with similar gene expression values to our putative new cell type [16–19]. The known functions of the highly expressed genes are analyzed, and we hypothesize the function of the unknown rare cell types [23, 24].

## Acknowledgments

This study began as Zoe Weiss’s summer Research Summer Institute project. She would like to thank Professor Guo-Cheng Yuan for supervising the beginning of this project. Zoe Weiss also thanks Professor Vicki Hertzberg for providing details about spatial statistics.

## 1 Declarations

### 1.0.1 Data Availability

This algorithm has been implemented in R and deposited at Github with URL https://github.com/zoeweiss/Rare-Cell-Type-Detection-Algorithm.git. The datasets analyzed in this study were obtained from the public domain.

